# The arms race in noncoding regions of transposable element: the evolution of anti-silencing and RNAi

**DOI:** 10.1101/2022.12.19.521048

**Authors:** Taku Sasaki, Kae Kato, Aoi Hosaka, Yu Fu, Atsushi Toyoda, Asao Fujiyama, Yoshiaki Tarutani, Tetsuji Kakutani

## Abstract

Transposable elements (TEs) are among the most dynamic parts of genomes. Since TEs are potentially deleterious, eukaryotes silence them through epigenetic mechanisms such as DNA methylation and RNAi. We have previously reported that Arabidopsis TEs, called *VANDALs*, counteract epigenetic silencing through a group of sequence-specific anti-silencing proteins, VANCs. VANC proteins bind to noncoding regions of specific *VANDAL* copies and induce a loss of silent chromatin marks. Sequence-specific anti-silencing allows these TEs to proliferate with minimum host damage. Here, we show that RNAi efficiently targets noncoding regions of *VANDAL* TEs to silence them *de novo*. Target motifs of VANC, in turn, evolved to escape RNAi. Escaping RNAi could be the primary event leading to the differentiation of sequence-specific anti-silencing systems. We propose that this selfish behaviour of TEs paradoxically could make them less harmful to the host.

## Introduction

Transposable elements (TEs) are major constituents of the large genomes of vertebrates and plants. As the movement of TEs is mutagenic and potentially deleterious to the host, most of TEs are silenced by epigenetic mechanisms, such as DNA methylation, histone modifications, and RNAi (Hosaka & Kakutani 2018). On the other hand, at least some TEs should occasionally be active on an evolutionary timescale, as they are very dynamic evolutionarily. Interestingly, population epigenomic analyses in the flowering plant *Arabidopsis thaliana* revealed variation in the epigenetic states of TEs among natural accessions (Kawakatsu et al. 2016; Quadrana et al. 2016), consistent with the view that TEs are occasionally activated in natural populations. Thus, an important question would be how TEs are activated and silenced in natural populations.

Arabidopsis serves as an ideal model organism to investigate the regulation of TEs, where precise TE sequences throughout the genome and trans-acting mutations affecting epigenetic modifications of TEs are available (Ito & Kakutani 2014; Quadrana et al. 2016). In Arabidopsis mutants defective in DNA methylation, diverse TEs are transcriptionally de-repressed and mobilized, demonstrating the importance of DNA methylation for the immobilization of these Arabidopsis TEs (Miura et al. 2001; Singer et al. 2001; Kato et al. 2003; Lippman et al. 2004; Tsukahara et al. 2009; Mirouze et al. 2009). Interestingly, TEs activated by the loss of DNA methylation in the mutant remain mobile even in the wild-type (WT) background (Kato et al. 2004). The inheritance of TE mobility correlated with the inheritance of DNA methylation status (Vongs et al. 1993; Kakutani et al. 1999; Kato et al. 2004). Nevertheless, recovery of DNA methylation is found in TEs with small RNA (Teixeira et al. 2009). In plants, small RNA is involved in the *de novo* DNA methylation pathway called RdDM (RNA-directed DNA methylation). Components of RdDM have been extensively studied in Arabidopsis (Matzke & Mosher 2013). Nonautonomous TE copies often induce the silencing of autonomous copies (Slotkin et al. 2005; Burgess et al. 2020; Sasaki et al. 2022), and it is assumed that this homology-based silencing is mediated by small RNA. However, evidence for the impact of RdDM on TE mobility in natural populations remains limited. More importantly, it remains largely unknown how heritable DNA methylation and silencing of TEs can be erased during evolution.

As a mechanism for the activation of TEs, we have previously shown that TEs named *VANDALs* can counteract epigenetic silencing. *VANDAL21* is one of the TEs silenced in WT plants, but it is mobilized when DNA methylation is lost (Tsukahara et al. 2009). An intriguing feature of *VANDAL21* and other *VANDAL* family TEs is that they encode proteins named VANCs, which have anti-silencing activity (Fu et al. 2013; Hosaka et al. 2017; Sasaki et al. 2022); expression of a VANC protein from a transgene induces loss of DNA methylation, transcriptional de-repression of the encoded genes, and mobilization of some of them (Fu et al. 2013; Hosaka et al. 2017; Sasaki et al. 2022). Thus, the anti-silencing activity of VANC proteins would counteract the genome defence by the host to silence TEs by DNA methylation.

Interestingly, the anti-silencing effects of VANC proteins were very sequence specific; expression of VANC21 protein induces loss of DNA methylation only in *VANDAL21* copies, while other sequences, including other *VANDAL* member copies, are unaffected (Fu et al. 2013). ChIP-seq analyses revealed that VANC21 protein is localized specifically in noncoding regions of *VANDAL21* copies, which are enriched in tandem repeats with specific short motifs (Hosaka et al. 2017). That short motif is also important for binding of VANC21 to DNA *in vitro*. A related *VANDAL* TE, called *VANDAL6*, encodes related proteins, and the expression of one protein, named VANC6, induces loss of DNA methylation in *VANDAL6* copies and related TE copies but not *VANDAL21* copies. The regions affected by VANC6 also have tandem repeats with motifs different from those in VANC21 targets. Thus, although *VANDAL21* and *VANDAL6* are similar in their overall sequences, the encoded VANC proteins and their target sequences are differentiated and have distinct specificities. This specific anti-silencing would allow the *VANDAL* TEs to proliferate with minimum damage to the host, which would be advantageous for the survival of the TEs within the host population, at least in the long term.

Here, we genetically characterized the ability of the host to re-silence the activated *VANDAL21* TEs. Genetic analyses revealed that RNAi targets DNA methylation in noncoding regions of activated TEs to immobilize them. Thus, both silencing and anti-silencing target noncoding regions of TEs. Furthermore, as small RNAs can function in trans to silence TEs, the rapid evolution of VANC targets could also be understood as an escape from RNAi.

## Results

### The VANDAL21 transgene activates endogenous VANDAL21, but the activated endogenous copy is re-silenced when the transgene is segregated

As is the case for most TEs in plant genomes, VANDAL family TEs are generally heavily methylated and silenced in WT plants. However, when WT plants are transformed with a full-length endogenous mobile copy of *VANDAL21*, called *Hiun* (*Hi*), endogenous *Hi* loses DNA methylation, is transcriptionally derepressed, and is mobilized (Fu et al. 2013; Appendix Figure S1, Figure 1). Interestingly, when the *Hi* transgene (*Hi*TG) was segregated in the self-pollinated progeny of the transgenic line, endogenous *Hi* was immobilized immediately (Fu et al. 2013; Fig. 1A), and the DNA methylation recovered almost to the original WT level (Fu et al. 2013; Fig. 1B). Analogous loss and recovery of DNA methylation can also be seen in other copies of *VANDAL21* (Fig. 1C). These results suggest that hosts have efficient mechanisms to methylate and silence *VANDAL21* in the absence of VANC21 protein.

**Figure 1.**
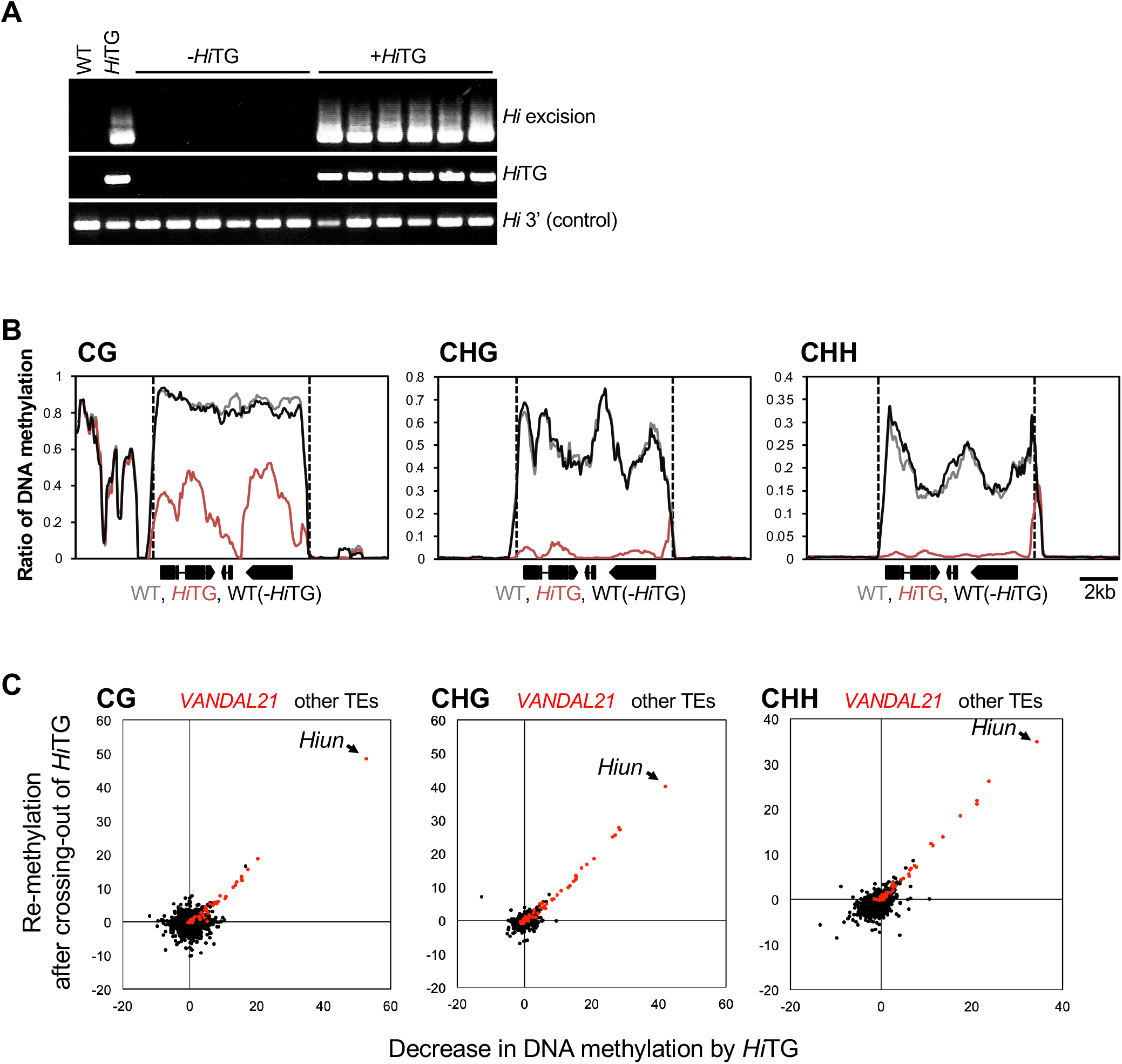
Endogenous *VANDLA21* TEs are re-silenced when *Hi*TG is segregated. (A) Excision of endogenous *Hi* in plants with and without *Hi*TG obtained from the same parental plant. (B) Patterns of DNA methylation for each cytosine context in an endogenous *Hi* locus calculated from WGBS data. (C) Comparison of the significance of changes in DNA methylation by *Hi*TG (x-axis) and those when *Hi*TG was segregated (y-axis) for all Arabidopsis TEs. The significance of changes in DNA methylation was assessed as described in Fu *et al*. (2013) (for details, see *Materials and Methods*). Dots in red and black indicate *VANDAL21* and other TEs, respectively.

### RNAi contributes to the re-silencing of Hi

To understand the underlying mechanism for this re-silencing, we examined the factors necessary for re-silencing. The Arabidopsis genome encodes three types of DNA methyltransferases for DNA methylation in CpG and non-CpG contexts (hereafter called mCG and mCH, respectively). MET1 maintains mCG, CMT2 and CMT3 directs mCH to regions with histone H3 lysine 9 methylation (H3K9me), and DRMs catalyse both the mCG and mCH *de novo* RdDM pathways (Lloyd and Lister 2022). In the *cmt2 cmt3* double mutant (hereafter called *cmt23*), *Hi* was re-silenced when *Hi*TG segregated (Fig. 2A). On the other hand, in the background of the *drm1 drm2* double mutant (hereafter called *drm12*), *Hi* often remained mobile even after the loss of *Hi*TG (Fig. 2B). Mutations in the other components of the RdDM pathway, such as NRPD1, NRPE2, NRPE1, and RDR2, also compromised the re-silencing (Appendix Fig. S2). Taken together, these results demonstrate that RdDM is effective for re-silencing *VANDAL21*.

**Figure 2.**
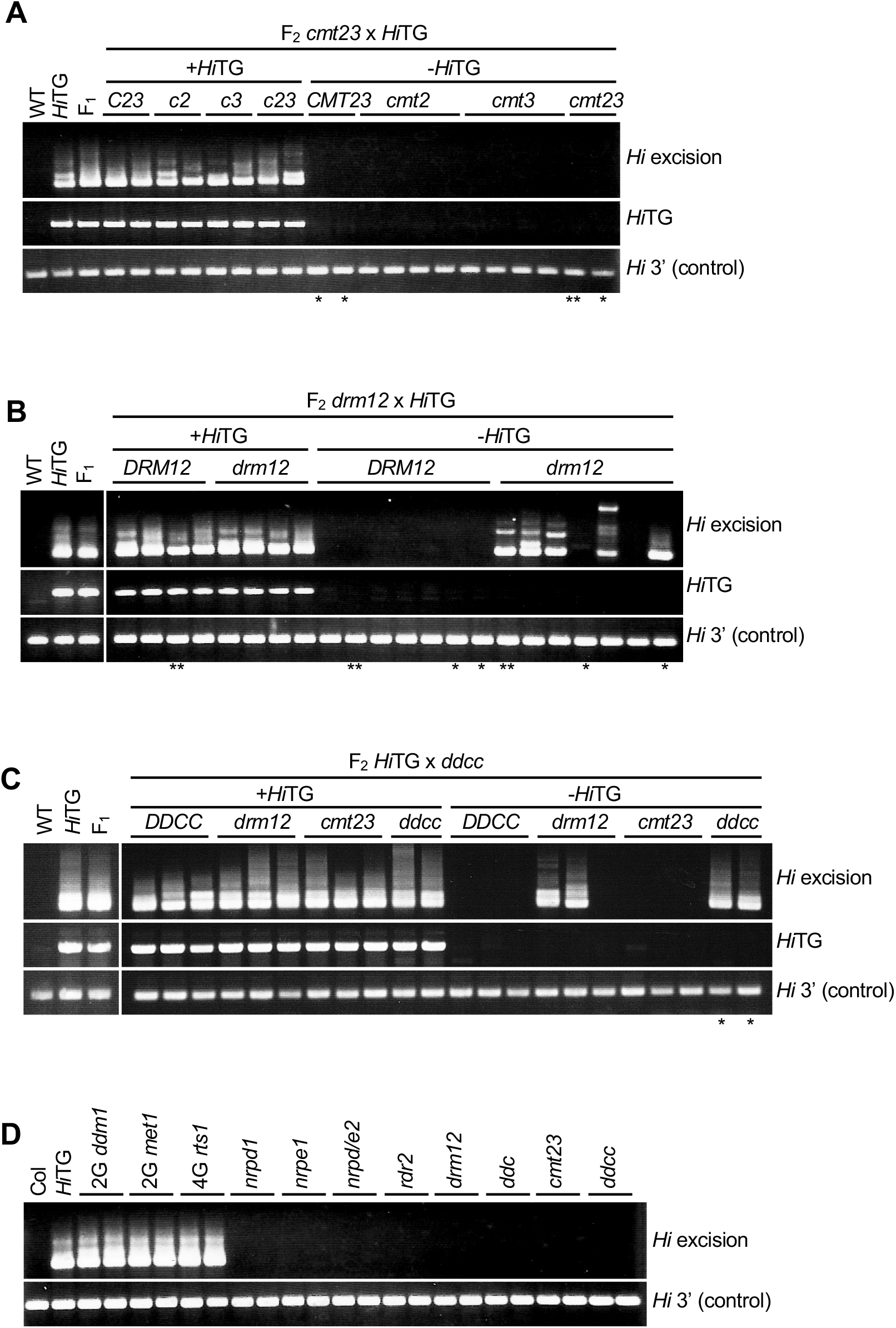
Re-silencing of *VANDAL21* depends on RdDM. (A-C) Excision of endogenous *Hi* in *cmt23* (A), *drm12* (B), and *ddcc* (C) mutant backgrounds when *Hi*TG is segregated. Plants with * and ** were used for WGBS, and the results of those with ** were used for the analysis shown in Figure 3 and Appendix Figure S5. (D) Excision of endogenous *Hi* in mutant backgrounds of epigenetic regulators. *drm12: drm1 drm2* double mutant, *ddc*: *drm1 drm2 cmt3* triple mutant, *cmt23*: *cmt2 cmt3* double mutant, and *ddcc*: *drm1 drm2 cmt2 cmt3* quadruple mutant.

Although *Hi* remained mobile when RdDM was compromised, we could not detect mobilization in a few sibling plants (two *drm12* plants shown in Fig. 2A; two *nrpd1*, one *nrpe1*, and two *nrpe2* plants shown in Appendix Fig. S2) in F2, as well as their self-pollinated F3 plants (Appendix Fig. S3), suggesting that additional mechanisms may also contribute to the immobilization of *Hi*.

A possible backup for RNAi-based DRMs in regard to mCH is CMTs; mCH is lost almost completely in the quadruple mutant *drm1 drm2 cmt2 cmt3* (hereafter called *ddcc*) (Stroud et al. 2014). Then, we analysed the immobilization of endogenous *Hi* in the *ddcc* mutant background. In the *ddcc* background, *Hi* remained mobile in all of the F2, F3, and F4 plants examined (Fig. 2C; Appendix Fig. S4), suggesting that CMTs are involved in the immobilization of *Hi* in the *drm12* mutant background. Importantly, without exposure to *Hi*TG, *ddcc* mutation does not induce the mobilization of endogenous *Hi* (Fig. 2D). Thus, *ddcc* mutation has a strong impact on the *de novo* immobilization of *Hi*, even though immobilized *Hi* remains immobile in the *ddcc* background.

The results above suggest that loss of mCH by *ddcc* does not affect the maintenance of immobilization. In striking contrast, *Hi* is mobilized when mCG is lost in the *met1* single mutant, demonstrating the critical role of mCG for the maintenance of *Hi* immobilization (Fig. 2D). *Hi* was also mobilized in *ddm1* and *rts1* mutants, which also abolished mCG in TEs (Fig. 2D). Taken together, these results suggest that mCG has a strong impact on controlling the activity of *VANDAL21*, while mCH machinery is necessary for *de novo* silencing of *VANDAL21*.

### RNAi is necessary for de novo CG methylation in targets of VANC21

The results above suggest the importance of mCG for controlling *VANDAL21*. In addition, the results demonstrate that RdDM is effective for *de novo* silencing of *VANDAL21* activated by VANC21. RdDM affects both mCG and mCH, and CH methylase CMTs can also contribute to *de novo* silencing. To further understand the role of mCG and mCH, we examined the impact of mutations in the RdDM machinery on the recovery of mCG and mCH. In the experiment for observing the recovery of DNA methylation after the loss of *Hi*TG, *drm12* mutation affected the DNA methylation recovery in noncoding regions more than in coding regions (Fig. 3A). These trends were also found in *VANDAL21* copies other than *Hi* (Fig. 3BC). Notably, ChIP-seq results suggest that VANC21 protein localizes in noncoding regions of *VANDAL21* copies (Hosaka et al. 2017). Indeed, DRM activity affected recovery around the peak of VANC21 target regions, which was defined by the loss of mCG (Fig. 4A). This effect was more conspicuous in mCG than in mCH (Fig. 4A). These results are consistent with the conclusion that mCG is important for *Hi* activity (Fig. 2D) and strongly suggest that RdDM silences *Hi* through *de novo* CG methylation of the target of VANC21.

**Figure 3.**
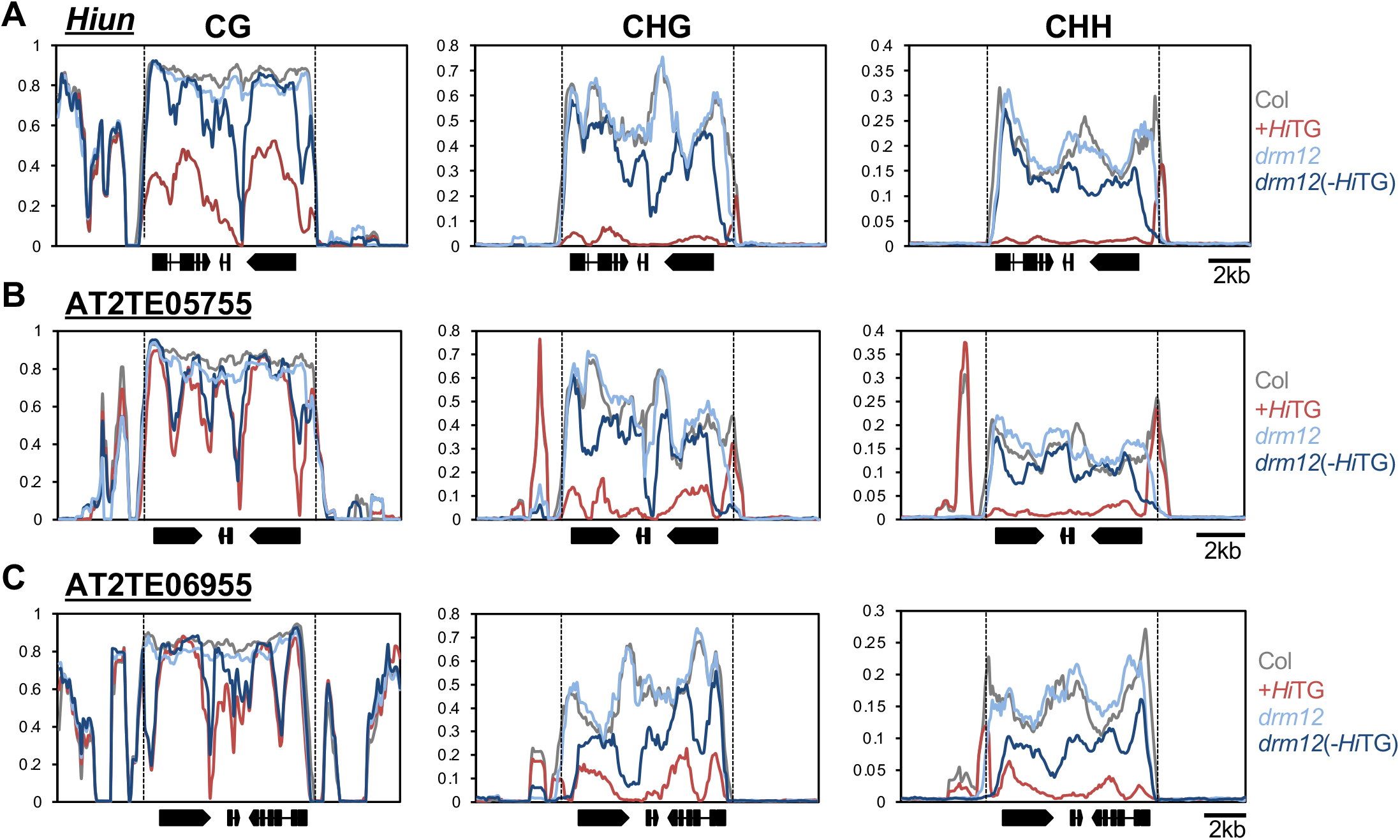
RdDM induces mCG in noncoding regions of *VANDAL21* TEs when *Hi*TG is segregated. (A-C) Patterns of DNA methylation in endogenous *Hi* (A), AT2TE05755 (B), and AT2TE06955 (C) in the *drm12* mutant background. ‘(-*Hi*TG)’ indicates that *Hi*TG was segregated. Dashed vertical lines indicate termini of TEs.

**Figure 4.**
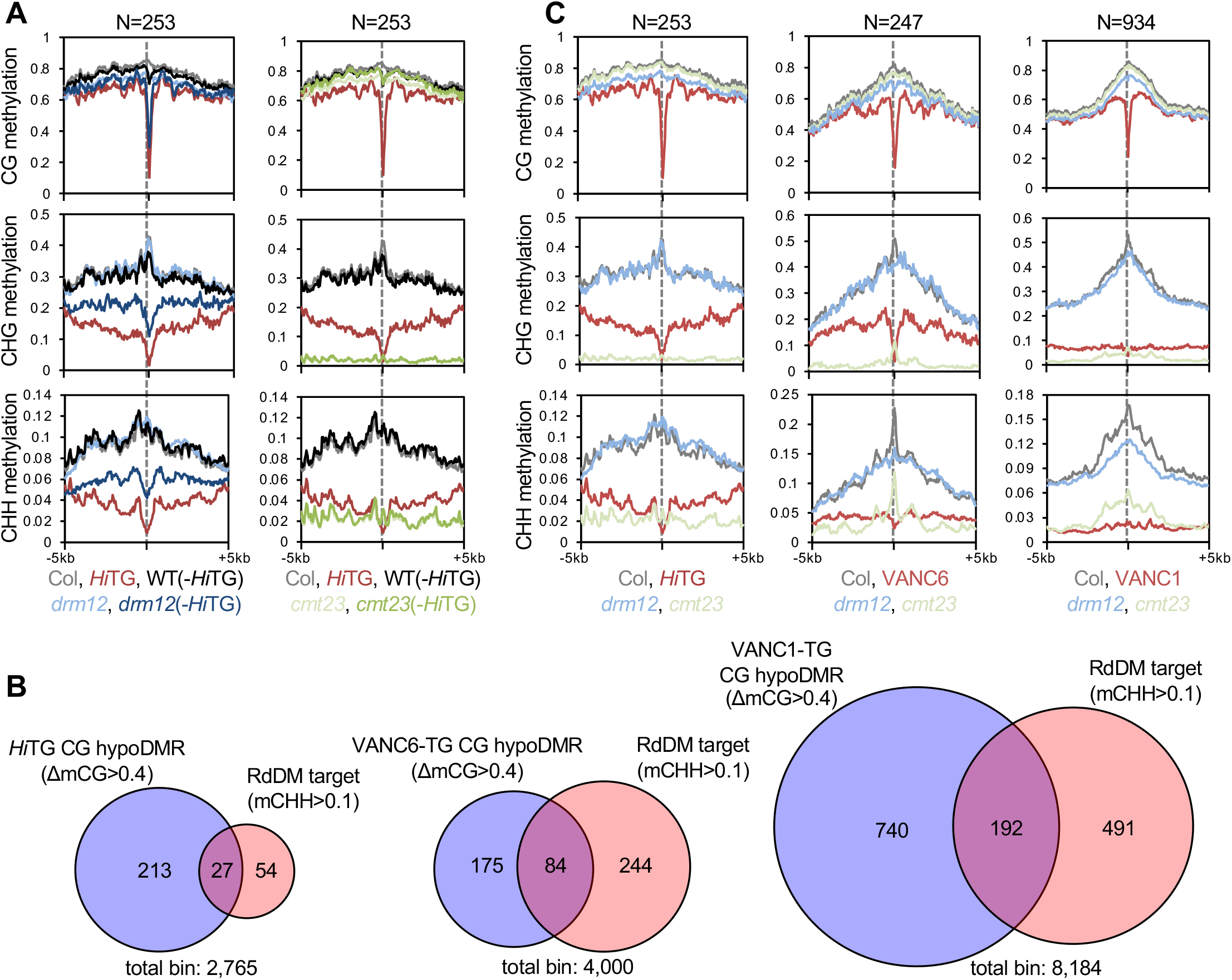
Target regions of RdDM and VANCs overlap. (A) Patterns of DNA methylation in flanking regions of VANC21 targets when *Hi*TG was segregated in *drm12* (left) and *cmt23* (right) mutant background. Grey dashed lines indicate the points of VANC targets. (B) Venn diagrams showing the overlap between targets of CG demethylation by VANCs and CHH methylation by RdDM. RdDM target regions are 3.13-, 3.96-, and 2.47-fold overrepresented in VANC21, VANC6, and VANC1 target regions, respectively (for VANC21, VANC6, and VANC1, hypergeometric *P*=1.880071666656578e-08, 5.417583973367242e-32, and 1.177936598724951e-36, respectively). (C) Patterns of DNA methylation around VANC21 (left), VANC6 (middle), and VANC1 (right) targets.

Complementary results were obtained for the effect of the *cmt23* mutations; in *cmt23*, mCG remained, although mCH was abolished nearly completely in coding regions (Appendix Fig. S5). Under such conditions, *Hi* was completely silenced and immobilized (Fig. 2A), again suggesting that mCG, rather than mCH, is critical for the maintenance of *Hi* silencing. Such complementary effects of RdDM and CMTs were also seen in other *VANDAL21* copies (Appendix Fig S5BC).

### Targets of RdDM and VANC overlap

The results above demonstrate that RdDM targets noncoding regions, as is the case for anti-silencing by VANC21. We therefore compared mCG DMRs hypomethylated in the presence of VANC21 (ΔmCG>0.4 in *Hi*TG) to RdDM targets (mCHH>0.1 in *cmt23*). RdDM targets were 3.13-fold overrepresented in VANC21-targeted regions (mCG DMRs) (hypergeometric p value = 1.880071666656578e-08) (Fig. 4B). Similar features were also observed in VANC1 and VANC6. Within the VANC1 and VANC6 target regions, RdDM targets were 2.47- and 3.96-fold overrepresented based on the same criterion (hypergeometric p value = 1.177936598724951e-36 and 5.417583973367242e-32, respectively, Fig. 4B). Moreover, target regions of VANC1 and VANC6 showed strong peaks of mCH in *cmt23* mutants, which reflects the activity of RdDM, although such a peak was less obvious in target regions of VANC21 (Figure 4C). Taken together, these results indicate that RdDM and VANC regulate common target regions in opposite directions.

### Differential CG methylation between HiTG and endogenous Hi

*Hi*TG induces reactivation of *VANDAL21*, which includes activation of the endogenous VANC21 gene (Appendix Fig. S1). Thus, an interesting question is why the activated endogenous VANC21 gene can be the target of re-silencing, while that of *Hi*TG can be expressed robustly. Endogenous *Hi* and *Hi*TG have the same sequences except for a few synonymous SNPs, but their responses to RdDM are different: while *Hi*TG activity is constant, endogenous *Hi* shows activity only when *Hi*TG is present, and *Hi* is targeted and silenced by RdDM when *Hi*TG is segregated. The differences in sensitivity to RdDM may reflect differences in the DNA methylation patterns. Indeed, bisulfite sequencing using locus-specific primers revealed that the DNA methylation status was different between endogenous *Hi* and *Hi*TG, while mCG remained in activated endogenous *Hi* (*Hiun*^*Hi*TG^), *HiTG* was free from mCG and mCH (Appendix Figure S6). Because their DNA sequences are nearly identical, differential mCG status is likely to be a determinant for sensitivity to RdDM.

To determine whether loss of mCG makes *Hi* tolerant to silencing by RdDM, we examined the behaviour of *Hi* activated by the *met1* mutation. The *met1-3* mutation induces a global loss of mCG and endogenous *Hi* is activated (Rigal *et al*. 2016, Figure 2D). We examined the behaviour of *met1*-activated *Hi* after introduction into the WT background. To distinguish between the origins of *Hi*, we utilized the GK-345D05 line for the WT parent; this line has a T-DNA insertion within the noncoding region of *Hi* (*Hiun*^GK345^) (Figure 5, Appendix Figure S7). We crossed the GK-345D05 line with the *met1-3* mutant, which has an endogenous *Hi* copy with mCG hypomethylation (*Hiun^met1^*), and generated the F2 population by self-pollination of F1 individuals. In the segregating F2 generation, we analysed the effects of *Hiun^met1^* and *Hiun*^GK345^ on the excision of endogenous *Hi*. While endogenous *Hi* was always active in the *met1* mutant background, the effect of *Hiun^met1^* and *Hiun*^GK345^ was monitored in the WT *MET1* background. Even in *MET1, Hiun^met1^* was actively excised, while most *Hiun*^GK345^ was re-silenced (Figure 5C). Consistent with their activity status, DNA methylation was restored in *Hiun*^GK345^, while *Hiun^met1^* remained hypomethylated despite intact RdDM machinery (Fig. 5D). These results indicate that complete loss of mCG makes *Hi* tolerant to silencing by RdDM.

**Figure 5.**
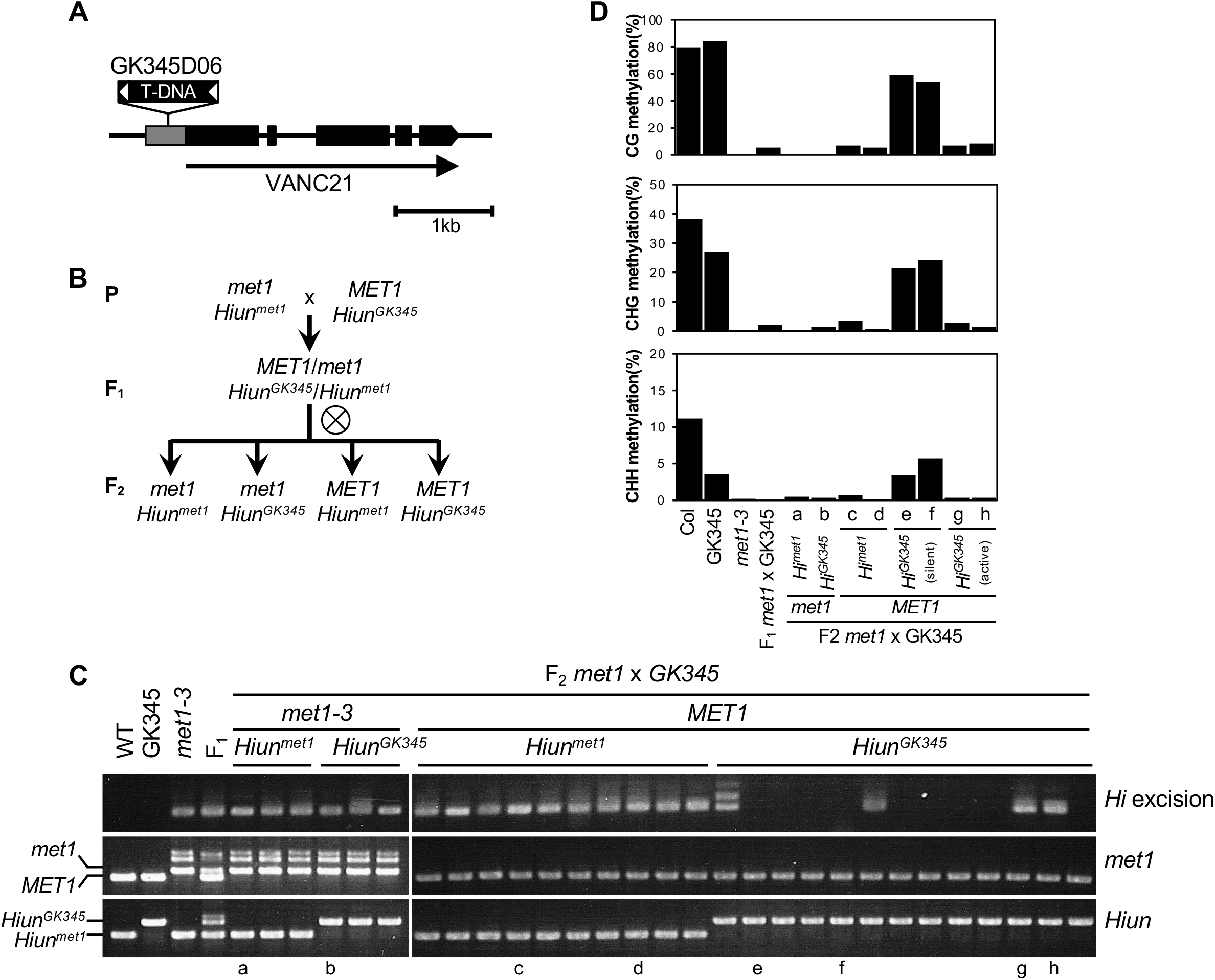
Loss of CG methylation makes *Hi* tolerant to RdDM-mediated silencing. (A) Schematic structure of GK345D06 (GK345), which has a T-DNA insertion within the noncoding region of *Hi*. (B) Genetic scheme to generate CG hypomethylated *Hi*. *met1* mutation induced a loss of CG methylation in *Hi* (*Hiun*^met1^), and GK345, which has *Hi* with DNA methylation similar to WT (*Hiun*^GK345^), enabled us to distinguish their origin. (C) *Hi* excision in F_2_ plants obtained from *met1* x GK345. Genotyping results of *MET1* and *Hi* loci are also shown. a-h indicate individuals who are analysed for their DNA methylation status in (D). (D) DNA methylation status in endogenous *Hi* calculated from the bisulfite sequencing results.

### CMT-mediated body mCH represses the transcription of VANDAL-encoded genes

TE genes generally have mCH in their internal regions (bodies), but the contribution of body mCH in TE silencing remains to be explored. While mCH in noncoding regions are targets of RdDM, body mCH are often catalysed by CMTs (To et al. 2020; To and Kakutani 2022). Consistent with this general trend, the body mCH of *VANDAL* TEs remained after the loss of the *Hi* transgene in the background of *drm12*. Interestingly, the mobility of *Hi* was more stably inherited in *ddcc* than in *drm12*. The WGBS revealed that the difference between *ddcc* and *drm12* is mainly in body mCH (Figure 6A). Consistent with the view that body mCH induces transcriptional repression, RT-qPCR revealed that the transcription of genes encoded within *Hi* was much higher in *ddcc* than in *drm12* (-*Hi*TG) (Fig. 6B). Interestingly, partial loss of body mCH in the *drm12 cmt2* (*ddc2*) and *drm12 cmt3 (ddc3*) mutant backgrounds resulted in partial activation. In addition, excision analysis of an endogenous *Hi* in the *ddc2* and *ddc3* mutant backgrounds also suggested that both CMT2 and CMT3 are required for effective immobilization of *Hi* (Appendix Fig. S8).

**Figure 6.**
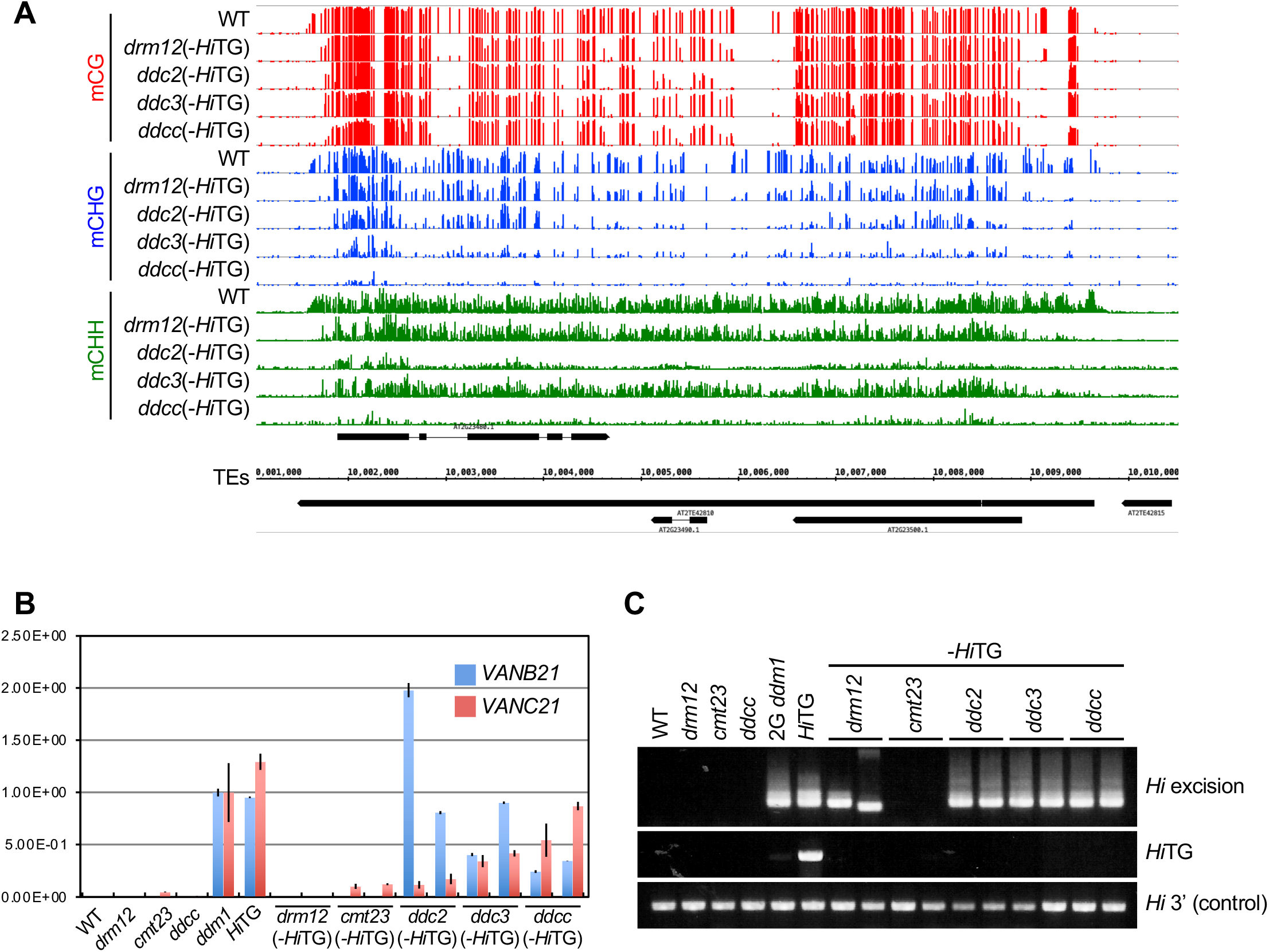
Gene-body mCH induces transcriptional repression of VANDAL-encoded genes. (A) Snapshot of DNA methylation status in the *Hi* locus in mutant backgrounds of DNA methyltransferases after *Hi*TG was segregated. (B) Expression of *VANB21* (At2g23490) and *VANC21* (At2g23480) measured by RT-qPCR. The values were normalized to the expression levels in the *ddm1* mutant. For each genotype, the average values and standard deviations of the three individuals are shown. Note that we could not detect the expression of VANC21 in *drm12* (-*Hi*TG), although endogenous *Hi* retained mobility (Fig. 6BC). This finding can likely be explained by the expression of VANA21, a transposase of VANDAL21, but we failed to compare the expression levels of this gene because of its very low expression level. (C) Excision of endogenous *Hi* in plants used for expression analysis in (B).

## Discussion

We have previously shown that the anti-silencing factor VANC21 binds to noncoding regions of *VANDAL21* TEs and induces loss of DNA methylation and mobilization (Hosaka et al. 2017). In this report, we characterized the mechanisms by which the host re-silences the activated *VANDAL21*: when the transgene expressing VANC21 is segregated, the RNAi-based *de novo* DNA methylation mechanism RdDM efficiently reintroduces DNA methylation in noncoding regions of the affected *VANDAL21* TEs and immobilizes them.

The binding regions of VANC proteins have specific motif sequences in tandem repeat organization. The tandem repeats contract and expand frequently and evolve synchronously. We have previously proposed that the synchronous evolution of short target motifs enables each of the VANDAL families to have different sequence specificities and proliferate with minimum host damage (Hosaka et al. 2017). On the other hand, tandem repeats can be sources of siRNA (Martienssen 2003) and targets of RdDM (Alleman et al. 2006). We found an overlap between the targets of RdDM and targets of VANC1/VANC6, although the overlap was less obvious for the targets of VANC21 (Fig. 4B). This unique feature of *VANDAL21* may reflect the fact that these elements are still proliferating and efficiently escaping RdDM. It is tempting to speculate that the short-term advantage of differentiation of the VANC systems for the TEs is the escape from RNAi. This short-term advantage of the evolution of the anti-silencing systems as an escape from RNAi would lead to their long-term advantage to maintain host fitness.

An intriguing feature of the effects of VANC on DNA methylation is that these effects spread outside the regions to which it binds and affect the entire TE. Although the effect is strongest in noncoding regions where VANC binds, surrounding coding regions also showed a significant loss of DNA methylation, especially in the CH context. Thus, anti-silencing by VANC seems to have at least two layers of effects: local effects on mCG and mCH in noncoding regions where it binds and additional effects, mainly for mCH, in surrounding regions (Figure 7). While the biological effect of VANC in noncoding regions seems significant, the role and control of VANC for hypomethylation in coding regions remain enigmatic. We showed that gene-body mCH contributes to the transcriptional repression of VANB21 and VANC21 (Fig. 6). Thus, the spreading of mCH loss has a significant impact on the stable maintenance of the anti-silencing effect. Interestingly, when the VANC transgene is segregated, loss of mCH in coding regions is recovered even in mutants of the RdDM machinery. Thus, in addition to the RNAi-based mechanism to silence VANDALs, RNAi-independent *de novo* silencing mechanisms may also function in coding regions. Consistent with this observation, RdDM-independent recovery of mCH of TE genes is generally observed after the introduction of the WT gene to mutants defective in mCH and H3K9me2 (To et al. 2020). In addition, *VANDAL21* activated by *met1* mutation remains active even in the *MET1* WT background. Thus, complete loss of mCG generates an allele tolerance to RdDM-based silencing; the effect of mCG on *de novo* establishment of mCH has also been detected in other TE genes (Chan et al. 2006; Sasaki et al. 2014; To et al. 2022). Crosstalk among multiple silencing pathways would achieve effective protection of the host genome from threat by TEs. The loss of DNA methylation in TEs by the transposon-encoded protein was also found for *Spm* (McClintock 1951, 1958). An *Spm*-encoded protein called TnpA has been shown to activate *Spm*, which correlates with DNA demethylation (Schläppi et al. 1994; Cui & Fedoroff 2002). Thus, anti-silencing by TEs may be more common than previously thought.

**Figure 7.**
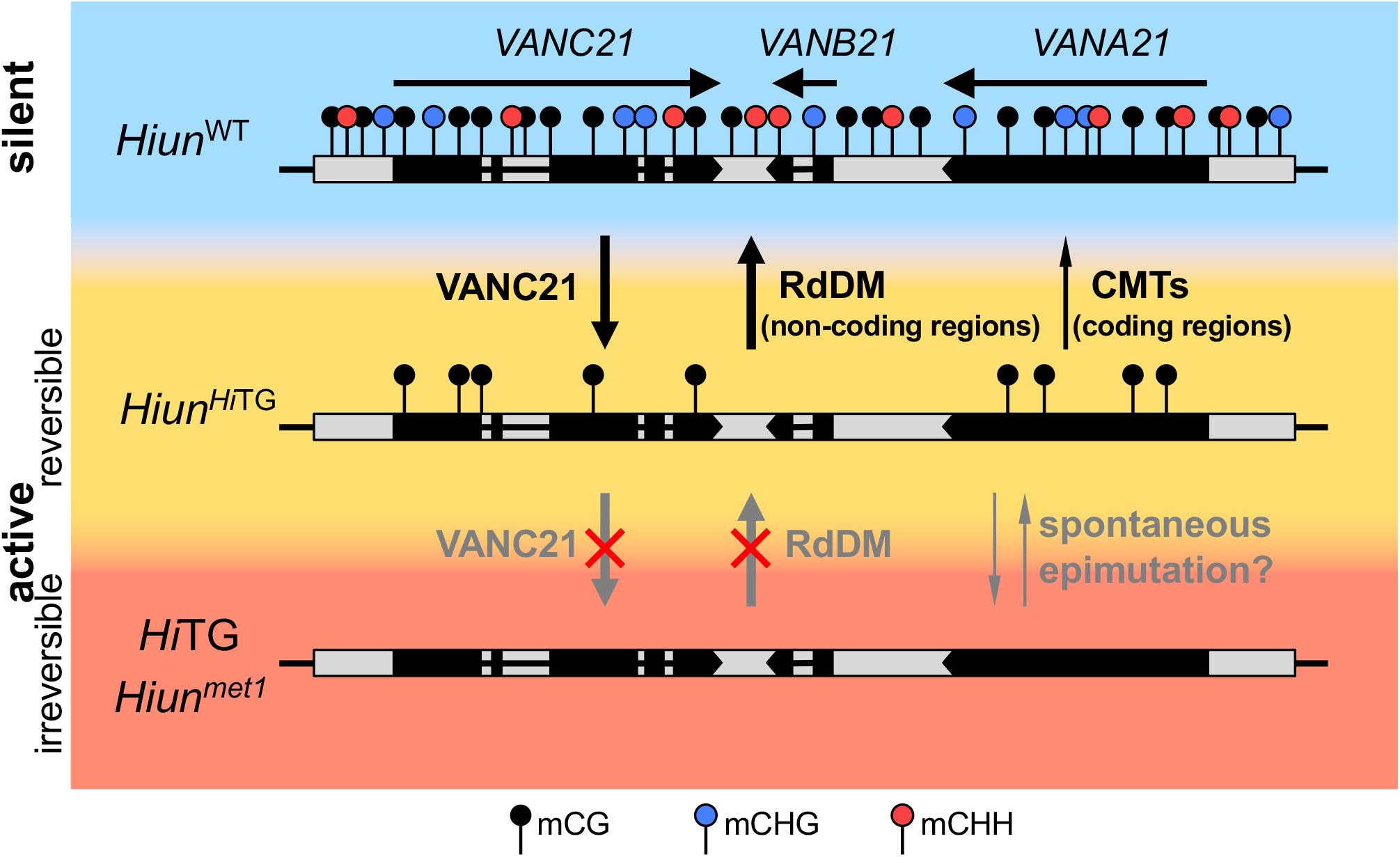
A possible model of the relationship between VANC21-mediated anti-silencing and RdDM. There are three different epigenetic states of *Hi*: constantly active (irreversible), active but reversible, and silent. *Hi*TG and *Hiun*^met1^ are constantly active, while endogenous *Hi* is active only when *Hi*TG co-exists (*Hiun*^*Hi*TG^) and reverts to a silent state (*Hiun*^WT^) when *Hi*TG is segregated. Among them, DNA methylation states are also different. *Hi*TG is free from DNA methylation, *Hiun^Hi^*^TG^ has mCG, and the silent *Hiun*^WT^ has both mCG and mCH. mCG has an important role in the recruitment of RdDM.

In this report, we showed that both VANCs and RdDM regulate noncoding regions of *VANDALs* as common targets in opposite directions. Another unique feature of *VANDAL* TEs is degenerated TIRs (terminal inverted repeats) (Le et al. 2000; Yu et al. 2000; Fu et al. 2013). The degeneration of TIRs can also be understood in terms of escape from RdDM; as long TIRs of *Mutator*-like TEs are often the source of siRNA (Burgess et al. 2020; Sasaki et al. 2022), degeneration of TIRs might compromise the pathway to induce *de novo* methylation of terminal regions of TEs because of low homology. Although a causative link remains to be tested, both degeneration of TIRs and the presence of VANC-related genes are commonly seen in VANDAL TEs. The arms race between RdDM and VANCs on the regulation of transpositional activity likely resulted in highly specific and divergent anti-silencing mechanisms of VANCs and contributed to the successful amplification of VANDAL TEs with minimal damage to the host.

## Materials and Methods

### Plant materials

Isolation of *met1-3, ddm1-1*, and *rts1-1/hda6-7* has been reported previously (Saze et al. 2003, Vongs et al. 1996, Aufsatz et al. 2002). The T-DNA insertion mutants *drm1-2* (SALK_031705), *drm2-2* (SALK_150863), *cmt2-3* (SALK_012874), *cmt3-11t* (SALK_148381), *rdr2-2* (SALK_059661), *nrpd1-3* (SALK_128428), *nrpe1-11* (SALK_029919), *nrpd/e2-2* (SALK_046208), and GK-345D05 were obtained from the Arabidopsis Biological Resource Center (Alonso et al. 2003, Kleinboelting et al. 2012). A transgenic line containing the full-length sequence of *Hi* (*Hi*TG) was reported by Fu et al. (2013).

### Detection of Hi excision

The excision of endogenous *Hi* was detected by nested PCR as described in Fu et al. (2013). Briefly, DNA was isolated from mature leaves using Nucleon Phytopure (GE Healthcare), and 10 ng DNA was used for nested PCR as a template. The PCR conditions for the first round of amplification were 94 °C for 1 min, followed by 25 cycles of 94 °C for 15 sec, 56 °C for 30 sec, and 72 °C for 45 sec, with a final extension at 72 °C for 3 min. PCR products were diluted 20 times with water, and 1 μl was used for the second round of PCR. The PCR conditions were the same as those for the first PCR except that the cycle number was changed to 30 cycles. The primers used for nested PCR are listed in Appendix Table S1.

### DNA methylation analysis

For bisulfite sequencing, 300 ng of genomic DNA isolated with Nucleon PhytoPure (GE Healthcare) was digested by *Pst* I in a 20 μl reaction mix. Digested DNA was ethanol-precipitated and eluted with 20 μl distilled water and then denatured by adding 2.2 μl 3 M NaOH and kept at 37 °C for 30 minutes. Denatured DNA was treated with 208 μl urea/bisulfite solution [7.5 g urea and 7.6 g sodium bisulfite dissolved in 20 ml water, adjusted to pH 5.0] and 12 μl 10 mM hydroquinone followed by 30 cycles of 95 °C for 30 sec and 55 °C for 15 min. Bisulfite-treated DNA was purified by a GeneClean III Kit (MP Biomedicals), eluted with 20 μl distilled water, and then desulfonated by adding 2.2 μl 3 M NaOH and incubating at 37 °C for 15 min. Finally, DNA was ethanol-precipitated and eluted with 20 μl distilled water. Converted DNA was used as a template for PCR using EpiTaq HS (TaKaRa). The PCR conditions were 94 °C for 1 min, followed by 35 cycles of 94 °C for 15 sec, 55 °C for 30 sec, and 72 °C for 1 min, with a final extension at 72 °C for 3 min. After gel electrophoresis, amplified DNA was purified and cloned into the pGEM T-Easy Vector System (Promega). At least 14 copies were sequenced for each sample. The primers used for this experiment are shown in Appendix Table S1.

For WGBS analysis, DNA extraction, bisulfite treatment, and library preparation were performed as described previously (Fu et al. 2013), and sequencing (paired-end, 150 bp) was performed by Macrogen Japan. Reads were trimmed using Trimmomatic (version 0.33) with the following parameters: “ILLUMINACLIP:TruSeq3-PE.fa:2:30:10 LEADING:3 TRAILING:3 SLIDINGWINDOW:4:15 MINLEN:36” (Bolger et al. 2014). Trimmed reads were mapped to the Arabidopsis genome (TAIR10) by Bismark ver. 0.15.0 (Krueger and Andrews 2011). The significance of changes in DNA methylation status was calculated as value (Mn/Cn - Mt/Ct)/(1✓Cn + 1✓Ct), where Mn, Cn, Mt, and Ct are methylated cytosine (M) and total cytosine (C) counts mapped for each TE in the nontransgenic (n) and transgenic (t) plants, respectively, as previously described (Fu et al. 2013).

### DMR detection

The DNA methylation ratio in each cytosine context within a 50 bp bin was calculated genome-wide, and regions whose CG methylation ratio changed more than 0.4 within each VANC target (*VANDAL21* for VANC21, *VANDAL6, 7, 8, 17*, and AT9TSD1 for VANC6, and *VANDAL1, 2, 22, 1N1*, and *2N1* for VANC1) were defined as DMRs. Because the DNA methylation status in TG is different from the endogenous status (Appendix Figure S4), bins overlapping with AT2TE42810 (*VANDAL21, Hiun), AT4TE25050 (VANDAL6*), and *AT1TE56425 (VANDAL1*) were excluded. For DMR detection in VANC6-TG and VANC1-TG, publicly available WGBS data (DRR099076 and DRR333229, respectively) were used (Hosaka et al. 2017; Sasaki et al. 2022).

### Hypergeometric test

For each VANC-target VANDAL family, TEs with >2 CG hypoDMRs were selected as VANC-target TEs. The DNA methylation ratio within a 50 bp bin for VANC-target TEs was calculated in Col, transgenic plants, and *cmt23* mutants.

### Expression analysis

Total RNA was isolated from mature leaves using TRIzol Reagent (Invitrogen) and treated with RQ1 RNase-free DNase (Promega). cDNA was synthesized using the PrimeScript III RT-PCR Kit (TaKaRa). Real-time qPCR (Figure 6A) was performed with Thunderbird SYBR qPCR Mix (TOYOBO) using 1 μl of cDNA diluted five times as a template. Three experimental replicates for each genotype were analysed. For RT-PCR (Appendix Fig S1B), 1 μl of 20-fold diluted cDNA was used for PCR as a template. PCR products were digested with *Pml* I, which digests only VANC21 transcribed from *Hi*TG. The primers used for RT-PCR are listed in Appendix Table S1.

## Data availability

The WGBS data obtained in this study are deposited in DDBJ under the accession number DRA014577.

## Author Contributions

TS and TK designed the project. TS, KK, AH, AT, AF, and YT performed the experiments. TS analysed the data. FY contributed materials. TS and TK wrote the manuscript.

## Acknowledgements

We thank Dr. Marjori Matzke for her comments on this manuscript. TS thanks Dr. Soichi Inagaki for the *ddc* and *cmt23* mutant seeds. This work was supported by grants from the Human Frontier Science Program (RGP0025/2021 to TK), CREST, Japan (JPMJCR15O1 to TK), the Japan Society for the Promotion of Science (JSPS) KAKENHI Grants (19H00995 and 21H04977 to TK, JP18K06348 to TS), and the National Institute of Genetics NIG-JOINT (32A2018 and 73A2019) to TS and YT. Computations for methylome analysis were performed on the NIG supercomputer at the ROIS National Institute of Genetics.

**Appendix Figure S1. Activation of endogenous *Hi* by *Hi*TG.**

(A) Schematic structure of *VANC21* and two synonymous mutations induced in *Hi*TG. These mutations created the recognition site of *Pml* I. Asterisks show the positions of mutations. Arrows indicate the positions of primers used for RT-PCR.

(B) Expression of endogenous and transgenic *Hi* detected by RT-PCR followed by restriction digestion by *Pml* I.

**Appendix Figure S2. Mutation of RdDM components compromises the re-silencing of endogenous *Hi* when *Hi*TG has segregated.**

(A-D) Excision of an endogenous *Hi* in mutants of RdDM components, such as *rdr2* (A), *nrpd1* (B), *nrpe1* (C), and *nrpd/*e*2* (D).

**Appendix Figure S3. RdDM-independent re-silencing was promoted in the F_3_ generation.**

(A-D) Re-silencing of endogenous *Hi* in mutants of RdDM components, such as *nrpd1* (A), *nrpe1* (B), *nrpd*/*e2* (C), and *rdr2* (D).

**Appendix Figure S4. RdDM-independent re-silencing for *Hi* activity was repressed in the *ddcc* mutant background.**

(A) Pedigree for materials used for analyses.

(B) Excision of endogenous *Hi* in the *ddcc* mutant background. (B) *ddcc* was fixed in the F_2_ generation, and excision in self-pollinated F3 plants with and without *Hi*TG was analysed.

(C) Excision of endogenous *Hi* in F_3_ *ddcc (-HiTG*) whose genotype was already fixed in F_2_.

**Appendix Figure S5. Patterns of DNA re-methylation in other *VANDAL21* TEs.**

(A-C) Patterns of DNA methylation are shown in Fig. 3 for *Hi* (A), AT2TE06955 (B), and AT2TE05755 (C) in the *drm12* and *cmt23* mutant backgrounds.

**Appendix Figure S6. Differential DNA methylation among each activity status.**

(A) Schematic structure of *Hiun*. Black boxes indicate exons. Positions of primers used for bisulfite sequencing are shown as black arrowheads.

(B-D) DNA methylation status of *Hi* in several genetic backgrounds detected by bisulfite sequencing.

**Appendix Figure S7. Importance of mCG for RdDM-mediated re-silencing of *Hi* (reciprocal cross of Fig. 5).**

(A) Genetic scheme to generate CG hypomethylated *Hi*. Individuals shown in red were used in this experiment, and those in black were used in experiments shown in Figure 4.

(B) Excision of endogenous *Hi* in the F2 population of WT x *met1* GK345. Note that *Hiun*^GK345^ retained mobility despite its T-DNA insertion.

**Appendix Figure S8. Gene-body mCH induces RdDM-independent re-silencing of *Hi*.**

(A, B) Excision of endogenous *Hi* in *ddc2* (A) and *ddc3* (B) mutant background. The pedigree for materials is shown in Appendix Figure S4A.

**Appendix Table S1.** Primers used in this study.

